# Uncovering the lung cancer mechanisms through the chromosome structural ensemble characteristics

**DOI:** 10.1101/2023.08.13.553145

**Authors:** Wen-Ting Chu, Jin Wang

## Abstract

Lung cancer is one of the most common cancers in human. However, it is still lack of understanding the mechanisms of a normal cell developing to the cancer cell. Here we develop the chromosome dynamic structural model and quantify the important characteristics of the chromosome structural ensemble of the normal lung cell and the lung cancer A549 cell. Our results demonstrate the essential relationship among the chromosome ensemble, the epigenetic marks, and the gene expressions, which suggests the linkage between chromosome structure and function. The analysis reveals that the lung cancer cell may have higher level of relative ensemble fluctuation as well as higher degree of the phase separation between the two compartments than the normal lung cells. In addition, the significant conformational *“*switching off*”* events (from compartment A to B) are more than the significant conformational *“*switching on*”* events during the lung cancerization. The kinetic lung cancerization pathway is not the same as the reversion pathway by characterizing the hot spots and interaction networks of the lung cancer transitions. These investigations have revealed the cell fate determination mechanism of the lung cancer process, which will be helpful for the further prevention and control of cancers.

## Introduction

Cancer is a group of diseases involving cancer cells with the potenial to invade or spead to other parts of the body while without normal grow, divide, and die cycle. Numerous studies have shown that mutations, genomic copy-number variations and translocations are common in cancer, which will lead to the de-regulation of gene expression ***Croce (2008)***. In addition, epigenetic alterations are also important for cancerization ***Baylin and Ohm (2006)***. Epigenetic alterations are functionally relevant modifications to the genome that do not change the nucleotide sequence, including DNA methylation, histone modifications, and changes in chromosomal architecture ***Kanwal and Gupta (2012); Timp and Feinberg (2013)***. *“*Epigenetic*”* was introduced by C.H. Waddington around 1940–1950 ***Van Speybroeck (2002)***. However, it is still challenging to understand how the epigenetic machinery affects the genomic structure as well as the gene expression.

In eukaryotic cells, three-dimensional (3D) chromatin structure is built up from the one-dimensional (1D) linear DNA sequence, correlated with the replication, transcription, and expression functions ***Jin et al***. (***2013); Cavalli and Misteli (2013); Bonev and Cavalli (2016); Rowley and Corces (2018)***. As the representative of chromosome conformation capture techniques, the recent developed Hi-C technique can capture 3D chromatin orgnization through high-throughput sequencing in a genome-wide fashion and the high-resolution chromosome pairwise contact map ***Lieberman-Aiden et al. (2009); Dixon et al. (2012); Rao et al. (2014); Stevens et al. (2017); Nagano et al. (2017); Tan et al. (2018); Bintu et al. (2018)***. Since 2009, many Hi-C results prove that the chromatin has certain critical *“*functional domains*”*, such as the topologically associating domains (TADs) and the compartments, which make up the hierarchically organized chromosome structure ***Lieberman-Aiden et al. (2009); Dixon et al. (2012); Rao et al. (2014); Hou et al. (2012)***. There are two different kinds of compartment types, A (euchromatin) and B (heterochromatin), which can be directly linked with the transcriptional activity (gene *“*switch on*”* or *“*switch off*”*) ***Lieberman-Aiden et al. (2009); Bickmore and van Steensel (2013); Solovei et al. (2016); Chu et al. (2022)***. In addition, the compartment A/B can reliably be estimated using epigenetic patterns ***Fortin and Hansen (2015)***. The TADs are local self-interacting domains identified within the compartments, which can regulate the gene expressions by restricting the promoters and enhancers within one TAD ***Dixon et al. (2012); Nagano et al. (2017); Gibcus et al. (2018); Chu and Wang (2020a***, 2021). TADs and compartments are the important structural and functional units of genome. Similar to proteins, the genome also has the *“*structure-function*”* relationship that different cell types (biological functions) alter in 3D chromatin structures. Therefore, the chromosome structure is the key for the investigation of the biological functions of genes and genomes. Besides, the chromatin immunoprecipitation (ChIP) technique with massive parallel sequencing method (ChIP-seq) was developed for identifying the genome-wide landscape of histone modifications ***Buck and Lieb (2004); Park (2009)***, determining the epigenetic patterns.

Cancer has a global significant impact on human organs, tissues, and cells, which will also affect the 3D structure of chromosomes, the hierarchical units, as well as the epigenomic profiles. A series investigations of Taberlay *et al*. have demonstrated that chromatin interactions may be important in the context of widespread, global genetic and epigenetic dysregulation and changes to the gene expression programs in carcinogenesis ***Taberlay et al. (2014***, 2016). Compartment switching (from A to B or from B to A) is observed at about 20% of loci level in the cancer cells ***Barutcu et al. (2015); Adeel et al. (2021); Yang et al. (2022)***. These significant compartment switching is associated with the gene expression changes in cancer cells. In addition, some essential changes on epigenetic markers (DNA methylation, histone modification patterns) have been discovered to influence the compartment states and the gene expressions, which may lead to the carcinogenesis. For example, the genes related to the histone acetyltransferase (HAT), histone deacetylase (HDAC), histone methyltransferases (HMT), DNA methyltransferases (DNMTs) enzymes are considered to contribute the epigenetic remodeling in cancer, including colorectal, bladder, non-small cell lung cancer, breast, prostate, and Wilms*’* tumor ***Audia and Campbell (2016); Bert et al. (2013); Chi et al. (2010); Khan et al. (2015)***. It remains to be further elucidated how the changes on the chromatin structure and epigenetics drive the gene expression differences and finally cause the shifts from normal cells to cancer cells.

The conformational space of a chromosomal structure is huge, given that Hi-C data capture the average chromatin behaviors of a population of cells. However, the Hi-C experiments have many limitations that they are costly to perform and the quality and reproducibility of Hi-C data are determined by the sampling cells ***Yardimci et al. (2019); Fiorillo et al. (2021)***. Therefore, certain important polymer models ***Rosa and Everaers (2008); Barbieri et al. (2012); Tjong et al. (2012); Le Treut et al. (2018); Annunziatella et al. (2018); Gürsoy et al. (2014***, 2017); ***Zhang and Wolynes (2015***, 2016); ***Di Pierro et al. (2016***, 2017); ***Liu et al. (2018a); Chiariello et al. (2016)*** have been developed for obtaining the 3D chromosome structures with hierarchies, territories, functional domains, as well as the ensemble behaviors, such as the model combines Minimal Chromatin Model (MiChroM) ***Di Pierro et al. (2016)*** and Maximum Entropy Genomic Annotation from Biomarkers Associated to Structural Ensembles (MEGABASE) ***Di Pierro et al. (2017)***. Here the MiChroM is a kind of heteropolymer model with different sub-compartment types of polymer beads. The MEGABASE establishes the relationship between the sub-compartment annotations and the histone modification marks from the ChIP-seq data. This type of chromosome model has been shown to successfully predict the 3D ensemble of chromosome structures of different cell types that are consistent with the experimental Hi-C data ***Di Pierro et al. (2017); Contessoto et al. (2021); Cheng et al. (2020); Junior et al. (2021); Contessoto et al. (2022)***. Perviously, we have developed the chromosome structural ensemble switching model (CSESM) based on MiChroM and MEGABASE, exploring the transdifferentiation pathways between PBMC and BN cell types on the Waddington*’*s landscape ***Chu et al. (2022)***. The results showed the fundamental relationship among chromosome structural ensemble behaviors, epigenetic patterns, and gene expressions. Aiming to uncover the mechanism of cancerization, here we focus on the dynamical chromosome structural changes between normal lung cell and the lung cancer cell (A549) to investigate the underlying *“*structure-function*”* relationship related to the lung cancer. By comparing the interactions, fluctuations, phases/compartments, and pathways of the cancerization process of several chromosomes, we observe the important impacts of cancerization on the chromosome structures as well as the functional domains. We further analyse the reversibility of the lung cancerization at different statistical levels. Finally, we show the gene loci involved in the significant compartment switching during the lung cancerization. Our results provide a quantitative and intuitive measure of the essential characteristics of chromosome structures and functions, implying the underlying mechanism of lung cancerization.

## Results and Discussions

### Two phases of chromatin have different behaviors in chromosomes

The Hi-C experiment can provide a contact probability matrix (*P*_*ij*_) to distinguish the euchromatin compartment A and the heterochromatin compartment B. Compartment B is more densely packed than compartment A while compartment A is more closely associated with open, accessible, actively transcribed chromatin. In the previous study, we have introduced a quantity called *“*local chromosome fluctuation index (local CFI)*”* 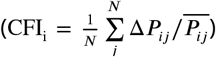, which has a strong correlation with the epigenetic quantity (compartment A/B) and gene expression ***Chu et al. (2022)***. That is, high local CFI values correspond to compartment A loci, which has high gene expressions. On the contrary, low local CFI values correlate with compartment B regions, which has relatively low gene expressions. Here in this study, we construct the ensemble for different chromosomes (17, 18, and 21) of normal lung and lung cancer (A549) cells. As the previous analysis, we quantify the epigenetic quantity, gene expression, and chromosome ensemble fluctuation with ChIP-seq data, RNA-seq data, and the local CFI value ***Chu et al. (2022)***. As shown in ***Figure 1***, there is a strong relationship among epigenetic quantity, gene expression, and chromosome ensemble fluctuation, which is consistent with our previous results. In addition, it is obvious that the chromosomes 17, 18, and 21 have different trends in the CFI vs. gene expression and local CFI vs epigenetic quantity figures (see ***Figure 1****a* and *c*). We notice that the local CFI has an important role in distinguishing the different chromosomes. The range of local CFI is about 10 to 16 in chromosome 21, which is significant lower than that in chromosomes 17 and 18. The differences between chromosome 21 and 22 (see ref ***Chu et al. (2022)***) are not as obvious as that among 17, 18, and 21. Therefore, the results suggest that the local CFI value is related to the chromosome length. The length of chromosomes 17 and 18 (about 83 and 80 Mb) is much longer than that of chromosomes 21 and 22 (about 47 and 51 Mb). The results also show that the chromosome fluctuation is linked to the chromosome length. This behavior can be explained by the decomposition results of the local CFI (see the micro CFI results in ***Figure 2)***. Long chromosome has additional long-range interactions between *i* and *j*, which will lead to larger value of the local CFI.

**Figure 1.**
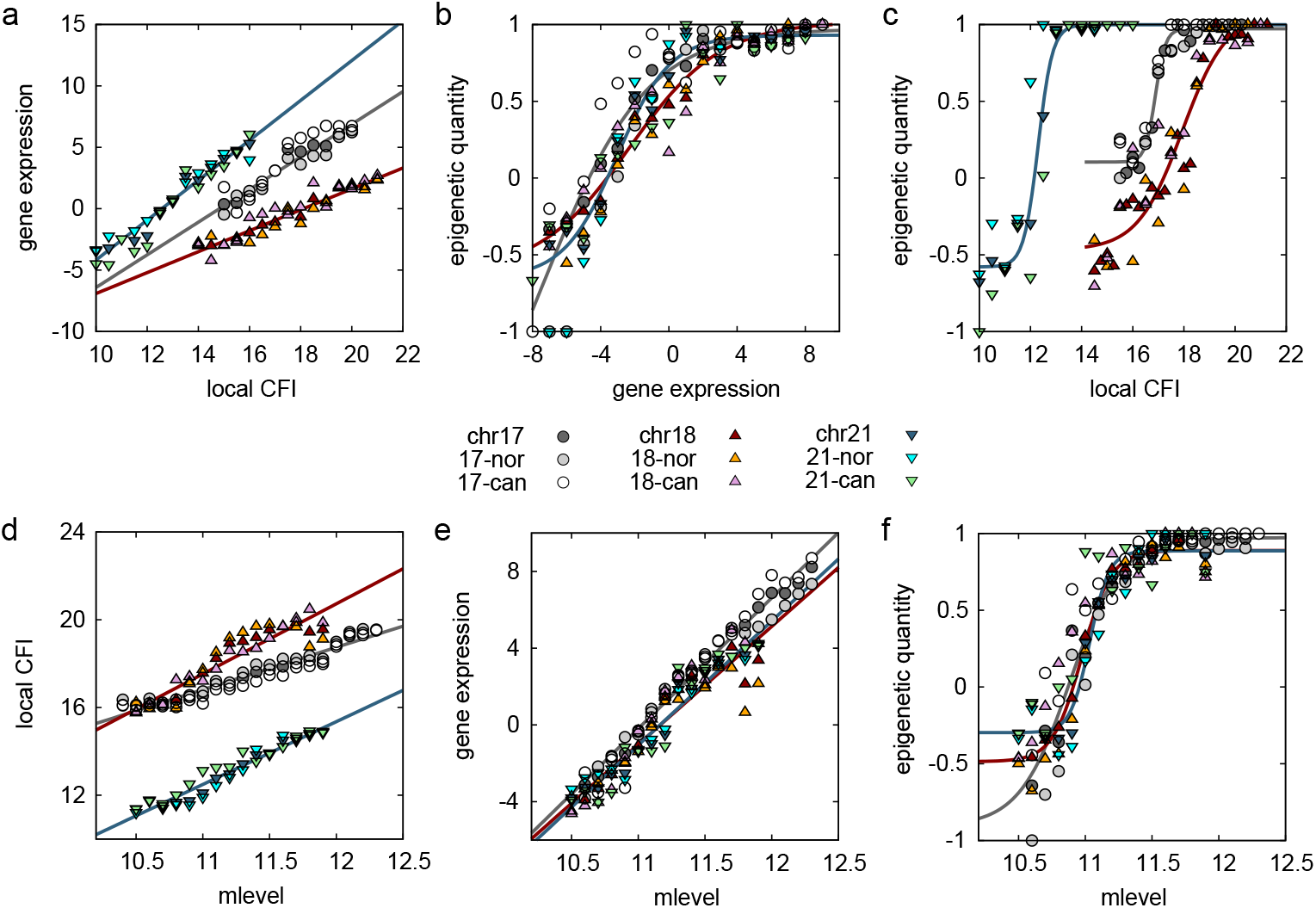
The correlation among epigenetic quantity, gene expression, chromosome ensemble, and DNA methylation level. (*a*) The distribution of the gene expression (calculated as ln(RNA-seq signal)) vs the local CFI and the fit lines; (*b*) the distribution of mean epigenetic quantity vs the gene expression and the fit lines; (*c*) the distribution of mean epigenetic quantity vs the local CFI and the fit lines; (*d*) The distribution of the DNA methylation level (calculated as ln(WGBS signal)) vs the local CFI and the fit lines; (*e*) the distribution of the methylation level vs the gene expression and the fit lines; (*J*) the distribution of the methylation level vs mean epigenetic quantity and the fit lines. Data of chromosome 17, 18, and 21 are illustrated with different colors and labels. Normal lung, lung cancer (A549), and the data of both, are shown in this figure.

**Figure 2.**
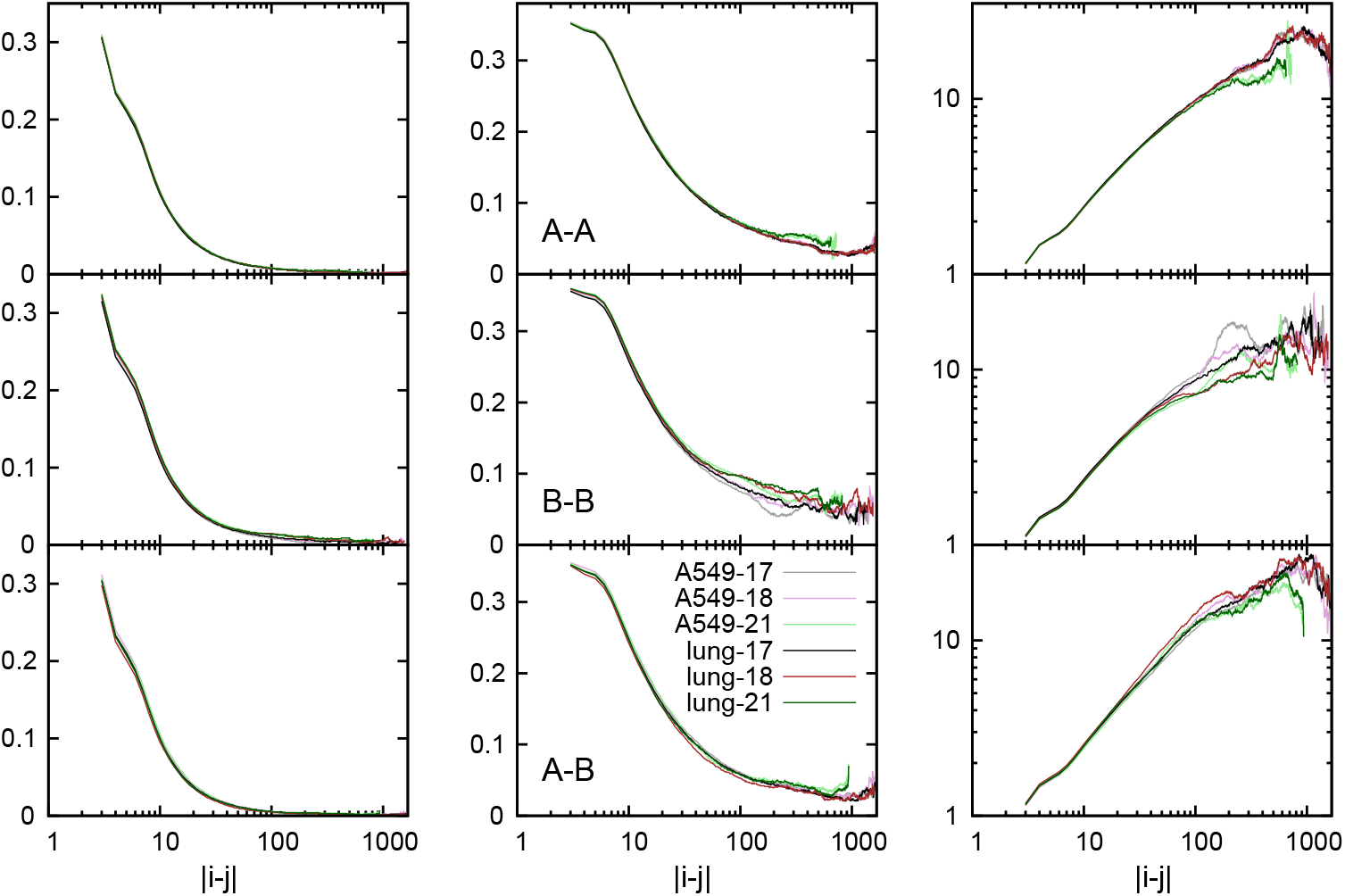
The mean contact probability 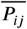 (left), the standard deviation Δ*P*_*ij*_ (middle), and the ratio of them 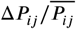 (micro CFI, right) within the compartment A (A–A), within the compartment B (B–B), and between compartment A and B (A–B) as a function of the chain distance |*i* − *j*| . Curves of normal lung and lung cancer (A549), chromosome 17, 18, and 21 are labeled in this figure.

Furthermore, we find out that the epigenetic quantity, gene expression, and chromosome ensemble fluctuation are all positively correlated with the methylation level (see ***Figure 1)***, which is quantified with the WGBS data (mean value per bead in 50 kb resolution). The results suggest that compartment A, which corresponds to high local CFI value, has higher methylation level than compartment B. This is consistent with the previous findings that high CpG island (CGI) density correlates with the compartment A and low CGI density correlates with the compartment B ***Liu et al. (2018b)***.

Though chromosomes 17 and 18 have similar length and local CFI range, at the same local CFI value, the level of gene expression and epigenetic quantity of chromosome 17 is a bit higher than that of chromosome 18. We believe that it has something to do with the ratio of compartment A/B (the number of beads belong to compartment A/the number of beads belong to compartment B). In chromosome 18, the ratio of compartment A/B is about 1.1 to 1.6 (*q* arm, lung and A549). While in chromosome 17, most loci are compartment A. The ratio of compartment A/B is about 4.8 to 5.2 (*q* arm, lung and A549). As a result, the ratio of compartment A/B may influence the distribution between the chromosome ensemble and the gene expression/epigenetic quantity.

### Cancerization has an impact on the chromosome phases and 2uctuation

The results above do not show the difference between normal lung and lung cancer chromosomes. Furthermore, we calculated the intra- and inter-compartment contact probability *P*_*ij*_ behaviors: mean value 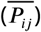, standard deviation (Δ*P*_*ij*_), and fluctuation 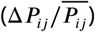 for each pair of loci *i* and *j*. According to the definition of local CFI (CFI_i_), we denote the individual fluctuation 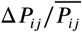 as microscopic-level CFI (micro CFI, CFI_ij_). As shown in Figure S1 and Figure S2, the compartment B is relatively stable (lower 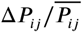 in B–B, with both higher 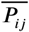 and Δ*P*_*ij*_). The trends of these data in A–A, B–B, A–B are consistent with the previous studies ***Chu et al. (2022)***.

Firstly we show the effect of chromosome length by comparing the data of chromosomes 17, 18, and 21 with the same loci interval. Overall, the effect of chromosome length is not obvious in the data of 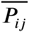, but is significant in the data of Δ*P*_*ij*_ and micro CFI (see ***Figure 2)***. In addition, the effect of chromosome length is more significant within the compartment B, than that within compartment A and that between compartment A and compartment B. The effect of the chromosome length is clearer for the long-range contact probability, especially the interval over than 100 beads (|*i* − *j*| ≥ 100). For different chromosomes, the results suggest that the shorter chromosome 21 has higher 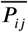 and higher Δ*P*_*ij*_ but lower micro CFI than those of the longer chromosomes 17 and 18.

Importantly, normal lung and lung cancer chromosomes can be distinguished in the figure of *P*_*ij*_ distribution (***Figure 2***, obviously in the figure of B–B). We find that the cancer cell A549 has lower 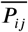 and lower Δ*P*_*ij*_ but higher micro CFI than the normal cell. We have shown in the previous study that the dimensionless quantity 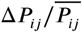 (micro CFI) is used to measure the relative fluctuation of the interactions in different compartments. Here the results indicate that the normal cell is relatively more stable (lower micro CFI) than the cancer cell. To show the differences of loci pairs between normal lung and A549 cells clearly, we calculated the contact probability *P*_*ij*_ (B–B, within the compartment B) behaviors during the pathway lung–A549 and the pathway A549–lung. As shown in ***Figure 3*** and Figure S3, normal lung and lung cancer chromosomes have significant differences in standard deviation and fluctuation, especially for the long-range contact (|*i* − *j*| ≥ 100). These results suggest that the cancerization (from normal lung to A549) will increase the relative fluctuation and cause the movement of the large chromosome domain.

**Figure 3.**
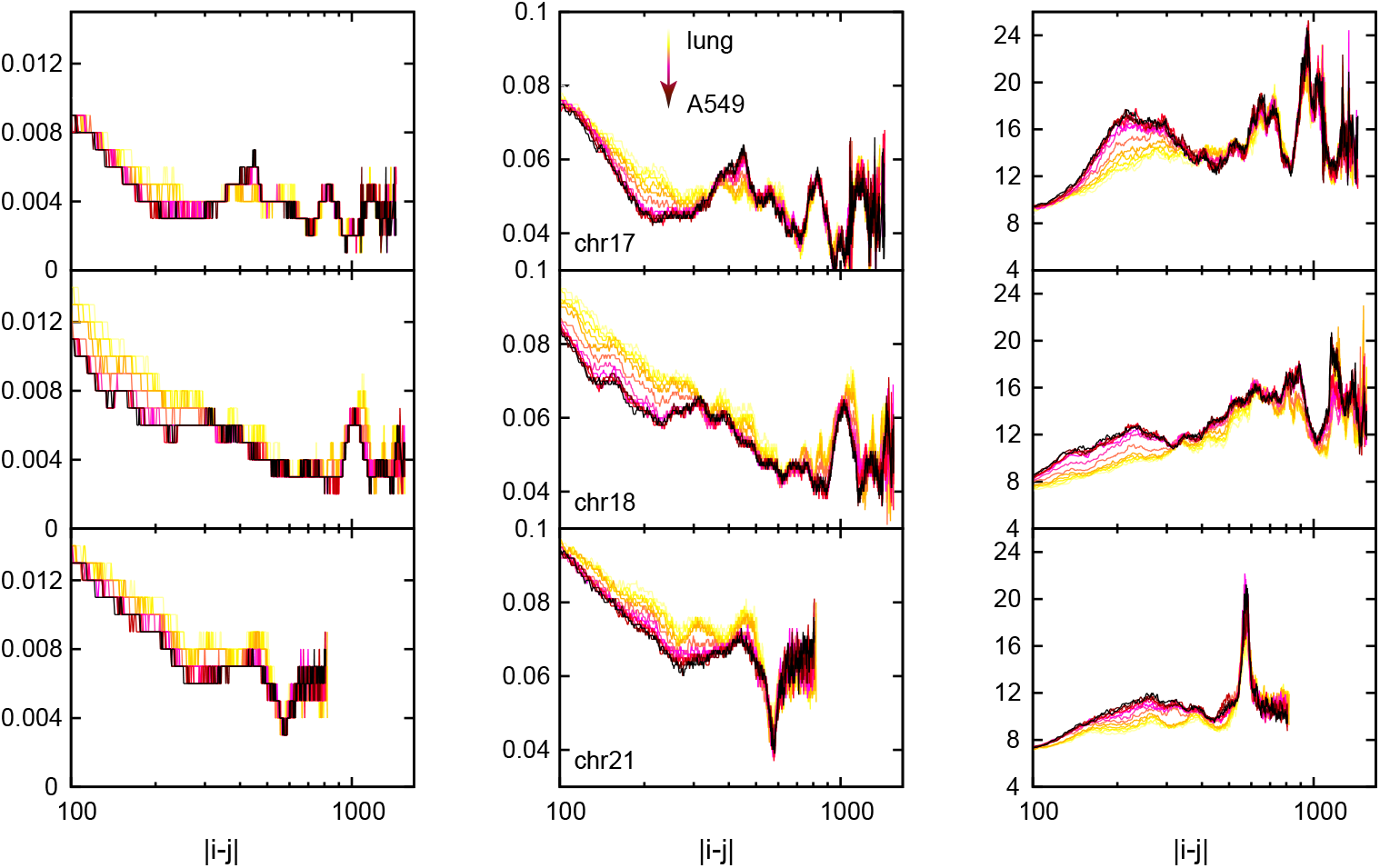
The mean contact probability 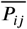 (left), the standard deviation Δ*P*_*ij*_ (middle), and the ratio of them 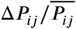 (micro CFI, right) within the compartment B (B–B) as a function of the chain distance | *i* − *j*| during the pathway from the normal lung cell to the lung cancer A549 cell. Curves of chromosome 17, 18, and 21 are labeled in this figure.

Moreover, we show the effect of cancerization on the distribution of phases and fluctuation. We set two parallel quantities to present the compartment reaction coordinate, *κ*_*A*_ (*α* ln(0.05−pc1_*i*_)(0.05− pc1_*j*_)) and *κ*_*B*_ (*α* ln(0.05 + pc1_*i*_)(0.05 + pc1_*j*_)). Here pc1 is the first principal component value that relates to the compartment type. Large positive *κ*_*A*_ values correspond to the interactions between *i* and *j* within compartment A (A–A); low positive *κ*_*A*_ values correspond to the interactions within compartment B (B–B); negative *κ*_*A*_ values correspond to the interactions between compartment A and compartment B (A–B). Similarly, large positive *κ*_*B*_ values represent the interactions within compartment B (B–B); low positive *κ*_*B*_ values represent the interactions within compartment A (A–A); negative *κ*_*B*_ values represent the interactions between compartment A and compartment B (A–B). For the quantity *κ* with similar range within compartment A (A–A) and within compartment B (B–B), both *κ*_*A*_ and *κ*_*B*_ were calculated for analysis. As shown in ***Figure 4***, Figure S4–S6, high *κ*_*A*_ (A–A) and high *κ*_*B*_ (B–B) values (≥20, both large pc1 values) correspond to high 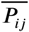 (close to 1), low Δ *P*_*ij*_ (close to 0), and low micro CFI (close to 0). These results suggest that these close and stable loci contacts (function regions) are located at the center of the phase (compartment A or compartment B). The *κ*_*A*_ and *κ*_*B*_ between 10 and 12 have moderately high 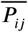 and moderately low micro CFI values. It seems that these loci contacts located at the periphery of the phase (compartment A or compartment B) may also have low loci intervals.

**Figure 4.**
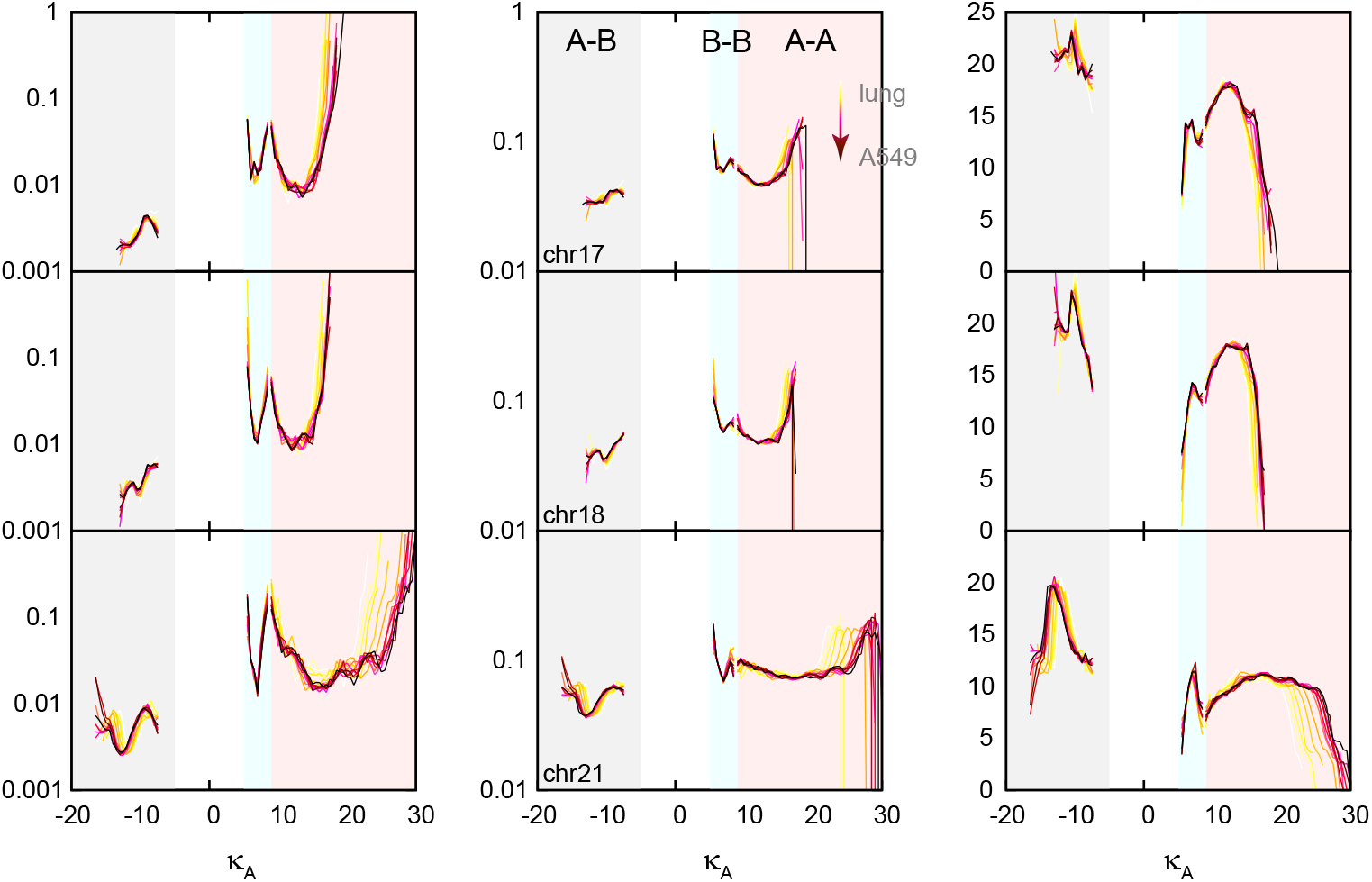
The distribution between *κ*_*A*_ (*α* ln(0.05 − pc1_*i*_)(0.05 − pc1_*j*_)) and 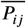 (left), between *κ*_*A*_ and Δ *P*_*ij*_ (middle), between *κ*_*A*_ and micro CFI during the pathway from the normal lung cell to the lung cancer A549 cell. The regions of different compartments, the curves of chromosome 17, 18, and 21 are labeled in this figure.

The distribution figures indicate that the cancer cell A549 has larger range of compartment than that of the normal lung cell. It is clear that the range of *κ*_*A*_ of A549 (about 10 to 30) is larger than that of normal lung (about 10 to 20) in chromosome 21 (see ***Figure 4*** and Figure S5). The results suggest that the cancer chromosome may have higher degree of the phase separation, for the first principal component (pc1) of the principal component analysis (PCA) represents the maximum variance direction in the data ***Abdi and Williams (2010)***. It may be resulted from the fusion of sub-compartments or TADs in the cancer chromosome. The previous studies have shown that the structural variation (SV) of chromosomes would result from *de novo* TAD formation, TAD-fusion events, and altered boundaries ***Spielmann et al. (2018); Wang et al. (2022)***. In addition, synergistic chromatin structure alteration at the TAD and compartment levels, for the insulation score changes as the compartments switch (from A to B or from B to A) ***Guo et al. (2021); Gridina and Fishman (2022)***. Therefore, the simulation results suggest that the TAD-fusion events is significant in the cancerization progress.

### Reversibility and irreversibility of the kinetic pathways

In this study, we performed the transition processes between normal lung cell and A549 (lung– A549 and A549–lung). Similar as the previous study ***Chu et al. (2022)***, we calculated the Pearson correlation coeffcient (*R*^2^) of the compartment (*8*) and local CFI (*γ*) spectrums to quantify the similarity between the two cells and show the overall transition path. In addition, the stem cell H1-hESC was added for comparison. As shown in ***Figure 5*** and ***Figure 6***, the overall forward and the reverse processes between lung and A549 seem to be reversible. There is no unified path between lung and A549 in different chromosomes, because the locations of lung, A549, and H1-hESC differ in chromosomes 17, 18, and 21. For the results of compartment, it is obvious that the length of path in chromosome 17 is shorter than those in chromosome 18 and chromosome 21. This may have something to do with the ratio of compartment A/B (see the analysis above). In chromosome 17, most loci are compartment A. For the results of local CFI, the locations of lung, A549, and H1-hESC in chromosome 17 and chromosome 18 are similar. We think it is because chromosome 17 and chromosome 18 have similar length (number of loci). The local CFI value can be linked with the chromosome length.

**Figure 5.**
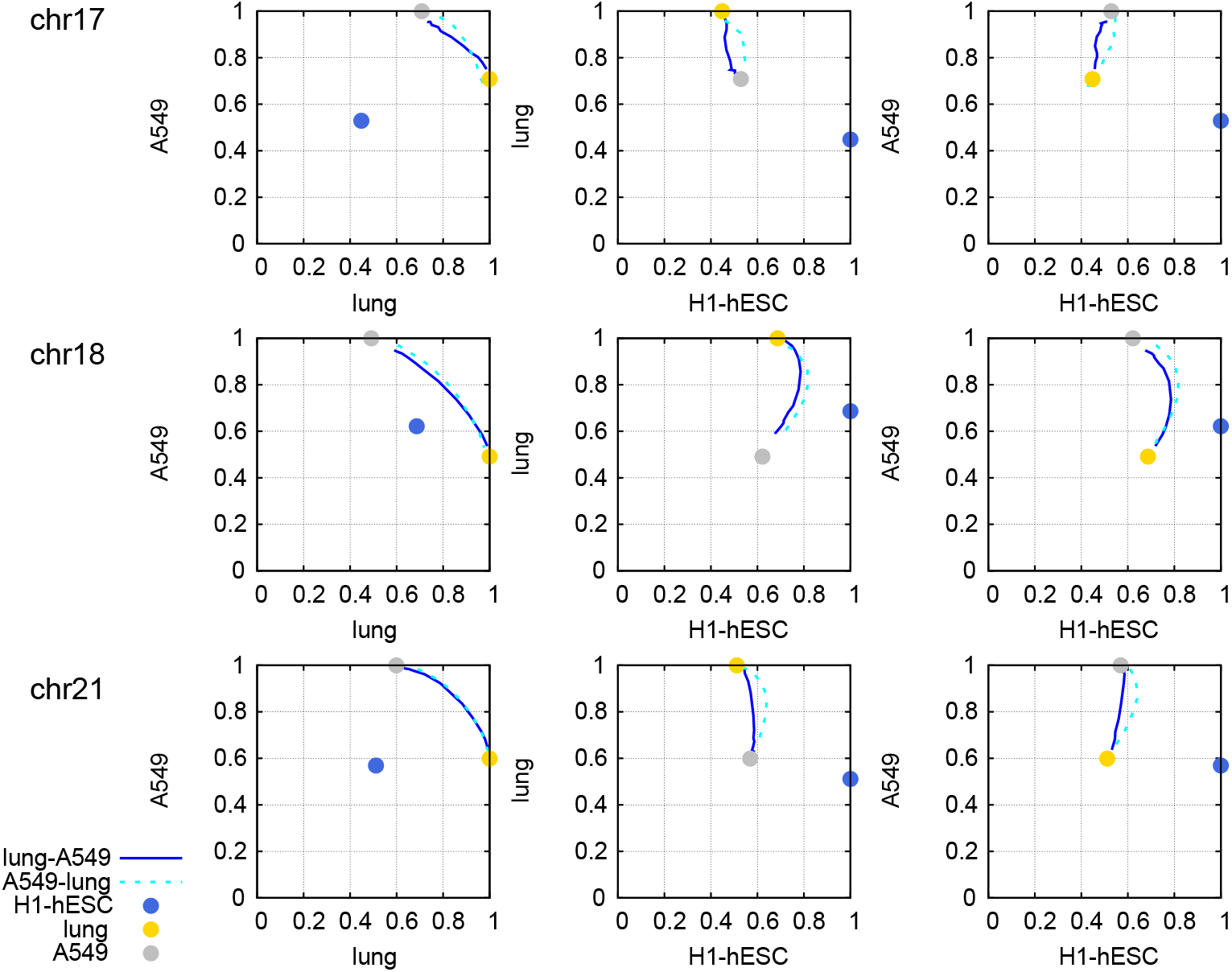
The kinetic pathways between normal lung cell and A549, including the forward processes lung–A549 and the reverse processes A549–lung, in chromosomes 17, 18, and 21. The trajectory is made up by the data at time 0*r* (start), 1*τ*, 3*τ*, 5*τ*, 10*τ*, 20*τ*, 50*τ*, 100*τ*, 150*τ*, 200*τ*, 250*τ*, 300*τ*, 350*τ*, 400*τ* (end). There are three reaction coordinates about compartment: *δ* _lung, *δ* _A549, and *δ* _H1-hESC. The locations of normal lung, A549, and H1-hESC cells are labeled.

**Figure 6.**
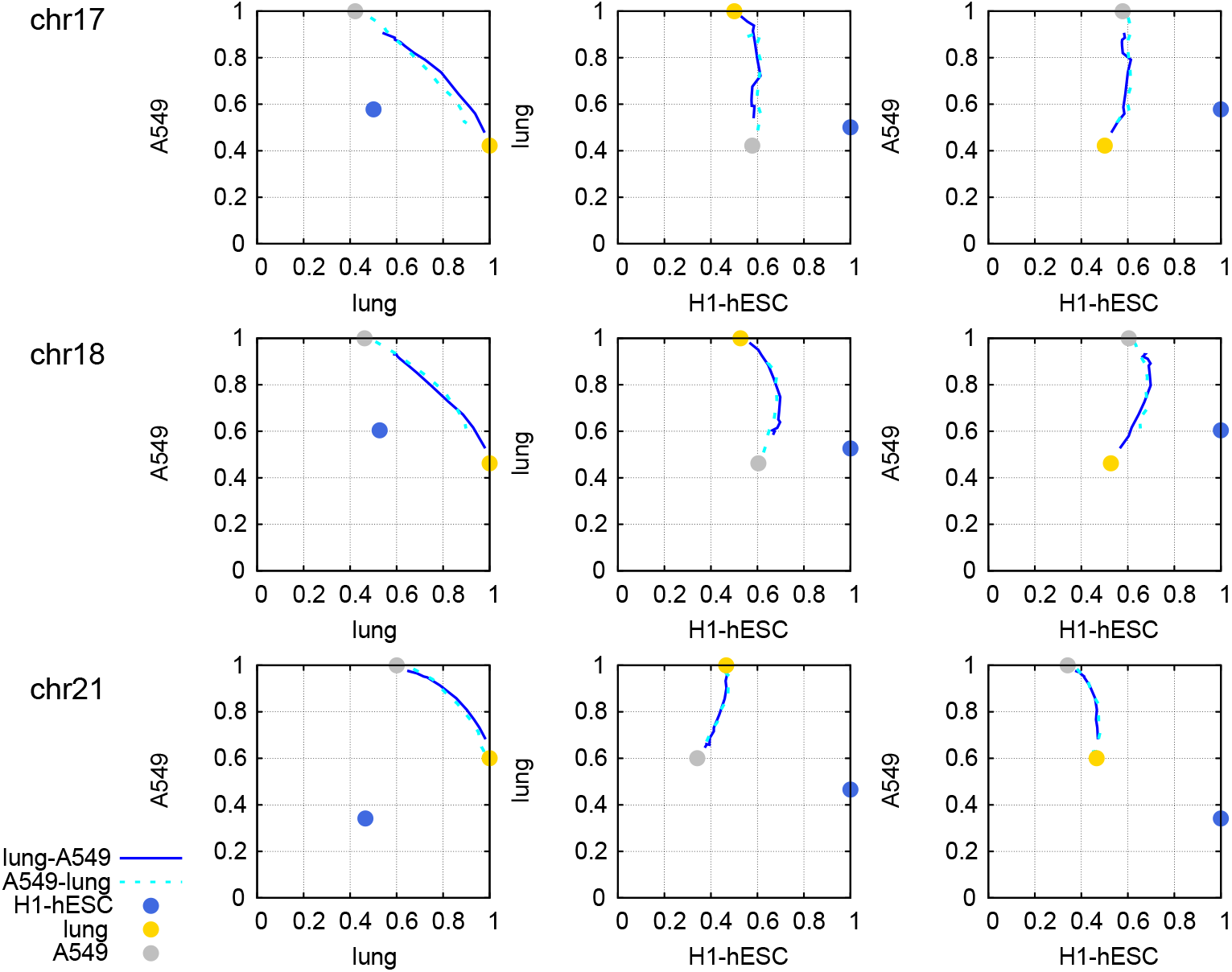
The kinetic pathways between normal lung cell and A549, including the forward processes lung–A549 and the reverse processes A549–lung, in chromosomes 17, 18, and 21. The trajectory is made up by the data at time 0*r* (start), 1*τ*, 3*τ*, 5*τ*, 10*τ*, 20*τ*, 50*τ*, 100*τ*, 150*τ*, 200*τ*, 250*τ*, 300*τ*, 350*τ*, 400*τ* (end). There are three reaction coordinates about CFI: *γ* _lung, *γ* _A549, and *γ* _H1-hESC. The locations of normal lung, A549, and H1-hESC cells are labeled.

In chromosomes, we have shown that the compartment A is linked with high transcriptional acitivity. When the compartment A is switched to the inactive region compartment B, the level of gene expression will decrease significantly. Therefore, we focus on the switching from compartment A to B (A–B) or from compartment B to A (B–A) during the cancerization (lung to A549) and the reversion (A549 to lung). The trajectories of the loci with significant A–B or B–A switching projected on both local CFI and compartment are illustrated in ***Figure 7**a*. The significant switching refers to the locus with obvious transition on both compartment and local CFI (compartment from A to B or B to A meanwhile the variation of local CFI larger than 1) during the cancerization and the reversion. Though there is no general A–B or B–A switching trend of the total loci, it is clear that much more significant *“*switching off*”* events than those significant *“*switching on*”* events occur during the cancerization in the chromosomes 17, 18, and 21. The ratio values of those significant A–B and B–A transition number are 4.3 (chr17), 11.0 (chr18), and 2.5 (chr21), respectively. The results suggest that the *“*switching off*”* (A–B) events may contribute significantly to the lung cancerization. And the range of local CFI changes of the *“*switching on*”* events is not as large as that of the *“*switching off*”* events. Similar trends can be found in the gene expression when we compare the RNA-seq data of the normal lung and lung cancer cells. As shown in ***Figure 7**b*, we define significant gene *“*switching off*”* and *“*switching on*”* loci by the difference of ln(RNA-seq signal) values. In the chromosomes 17, 18, and 21, the number of gene *“*switching off*”* loci is more than that of gene *“*switching on*”* loci, which is consistent with the simulation results of the conformational switching (compartment and local CFI). Furthermore, we have checked the RNA-seq data of other chromosomes and found the same trend of the genes after the lung cancerization (see Figure S7). Therefore, the gene *“*switching off*”* is more prominent than gene *“*switching on*”* for the lung cancer process.

**Figure 7.**
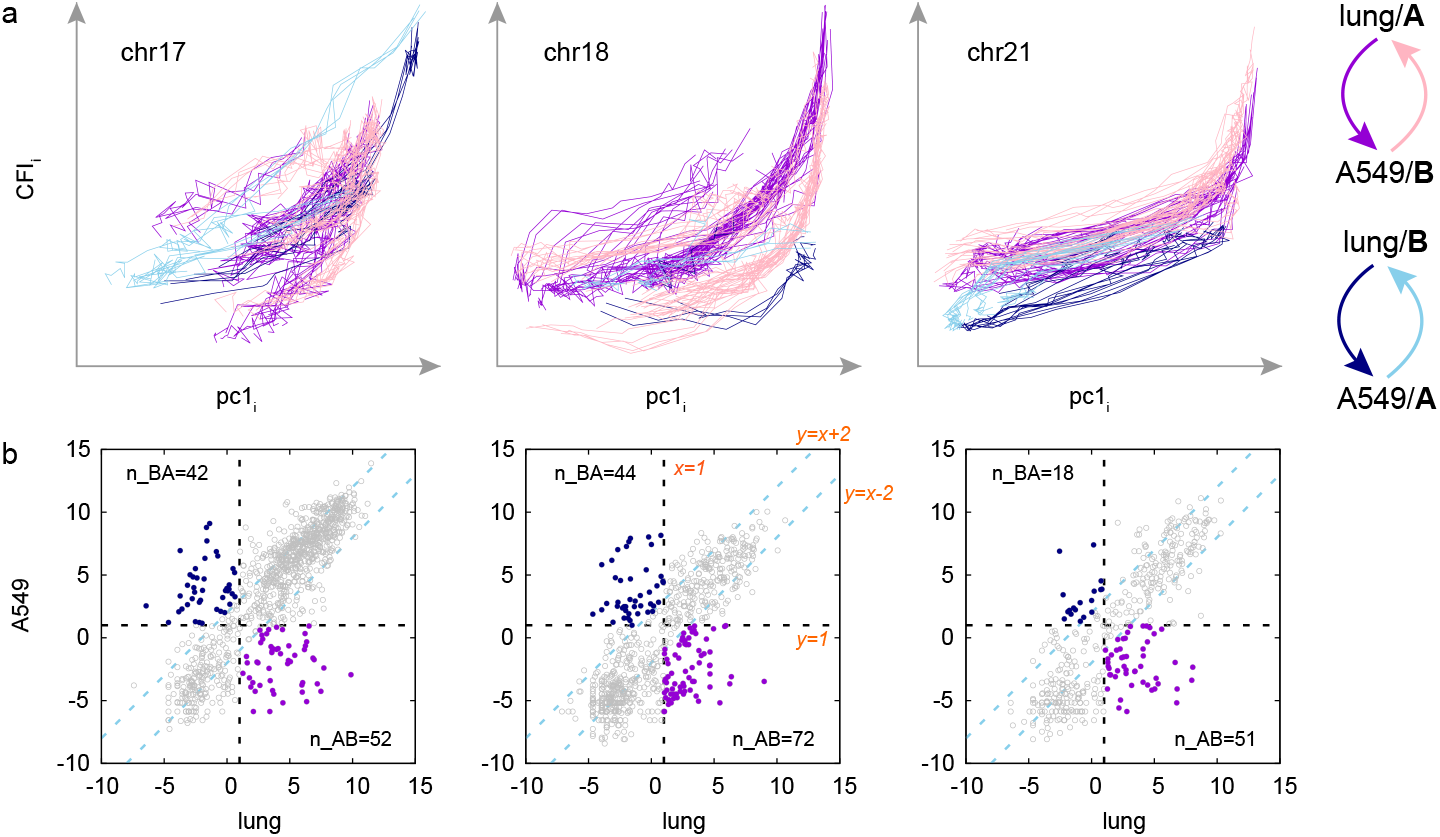
Significant switches of compartments and genes. (*a*) The distribution of local CFI (CFI_*i*_) and the compartment (pc1_*i*_) of the significant A–B or B–A switch during the forward processes lung–A549 and the reverse processes A549–lung; (*b*) The different gene expression level (quantified with ln(RNA-seq signal)) of each locus (in 50 kb resolution, *q* arm) in the normal lung and the lung cancer (A549) cells. In panel *a*, each line represents the trajectory of a locus (*i*) with significant A–B or B–A switch. Different processes and different type of switchs are illustrated with different colors. In panel *b*, the significant gene *“*switching off*”* (A–B, purple dots) or *“*switching on*”* (B–A, blue dots) loci during the cancerization process are defined with several lines, including *y* = *x* + 2, *y* = *x* − 2, *x* = 1, and *y* = 1. Other loci are shown as gray circles. The number of gene *“*switching off*”* (n_AB) and *“*switching on*”* (n_BA) loci is labeled in this panel.

The results above have already shown that the high DNA methylation level corresponds to compartment A, high gene expression level, and high local CFI value (***Figure 1)***. As a result, the significant *“*switching off*”* events (A–B) during the lung cancerization process from lung to A549 suggest that it is a hypomethylation process. Previous reports indicated that DNA repeats and large regions of DNA display extensive hypomethylation in cancer ***Feinberg and Vogelstein (1983); Esteller (2008)***. Therefore, global DNA hypomethylation is observed in cancer, which agrees with our results.

In addition, we can find out from the local CFI-compartment figure that the cancerization process cannot overlap with the reversion processes, which is obvious for the B–A transitions of the cancerization (***Figure 7****a*). If we separate the trajectory into each locus and project it into multiple reaction coordinates (local CFI and compartment), the path of lung–A549 and the path of A549–lung will be different (see Figure S8 as an example of three loci). Therefore, the cancerization pathway would be irreversible in multi-dimensional space.

### Kinetic hot spots and networks of lung cancerization

In proteins, Fersht *et al*. have uncovered the residues important for the protein folding kinetic stability by calculating the Φ value ***Fersht (1995); Clementi et al. (2000); Levy et al. (2005)***. Though some of these residues are not located at the active site (for binding), they can influence the kinetic barrier and mechanism when mutated to other residues. The Φ value is calculated as the ratio of free energy (fraction of contact probability) changes from transition state to unfolded state and from folded state to unfolded state. Here in this study, we plan to quantify the loci that have an impact on the kinetic stability of lung cancerization or reversion by calculating the Φ_*i*_ value with the local CFI changes: 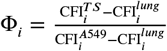 (cancerization) and 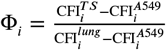 (reversion). We set the transition state (TS) as the conformation ensemble at kinetic time 50*τ* for the variation of the local CFI at this time is almost higher than that at other times. High Φ_*i*_ value refers to the locus changing first and crucial for the transition state formation. In order to avoid the abnormal Φ_*i*_ values (extreme high or low value because of very close 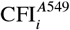 and 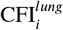 values), we do not caculate the Φ with 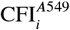 and 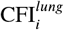 lower than 0.8.

The regions with high Φ values are illustrated in ***Figure 8*** (*a* cancerization and *b* reversion). These regions are important for the lung cancerization or reversion kinetic process in chromosomes 17, 18, and 21, including both protein-coding regions (genes) and non-coding regions. It is obvious that the high Φ loci of the cancerization pathway are different with those of the reversion pathway, which suggest the irreversibility of the cancerization and reversion pathways. Furthermore, changes in gene–gene interaction (gene network) has been reported to occur during the onset and progression of cancers ***Mair et al. (2019); Arshad and McDonald (2021)***. Aiming to find out the changes in gene– gene interaction during the lung cancerization or reversion, we quantify the gene–gene (locus–locus) correlation through the Pearson correlation coeffcient (*R*^2^) between the two traces of local CFI of bead *i* and bead *j* (here we use the data at time 0*τ* (start), 1*τ*, 3*τ*, 5*τ*, 10*τ*, 20*τ*, 50*τ*, 100*τ*, 150*τ*, 200*τ*, 250*τ*, 300*τ*, 350*τ*, 400*τ* (end) as the kinetic trajectory). The cross correlation matrix is illustrated in ***Figure 9***. High *R*^2^ value (close to 1) refers to high correlation between bead *i* and bead *j* that associated with the lung cancerization or reversion. The correlation network can be quantified with the pc1 of PCA results (see ***Figure 9***, blue lines). In addition, the network components of the cancerization pathway is not the same as those of the reversion pathway, which is consistent with the irreversibility conclusion above.

**Figure 8.**
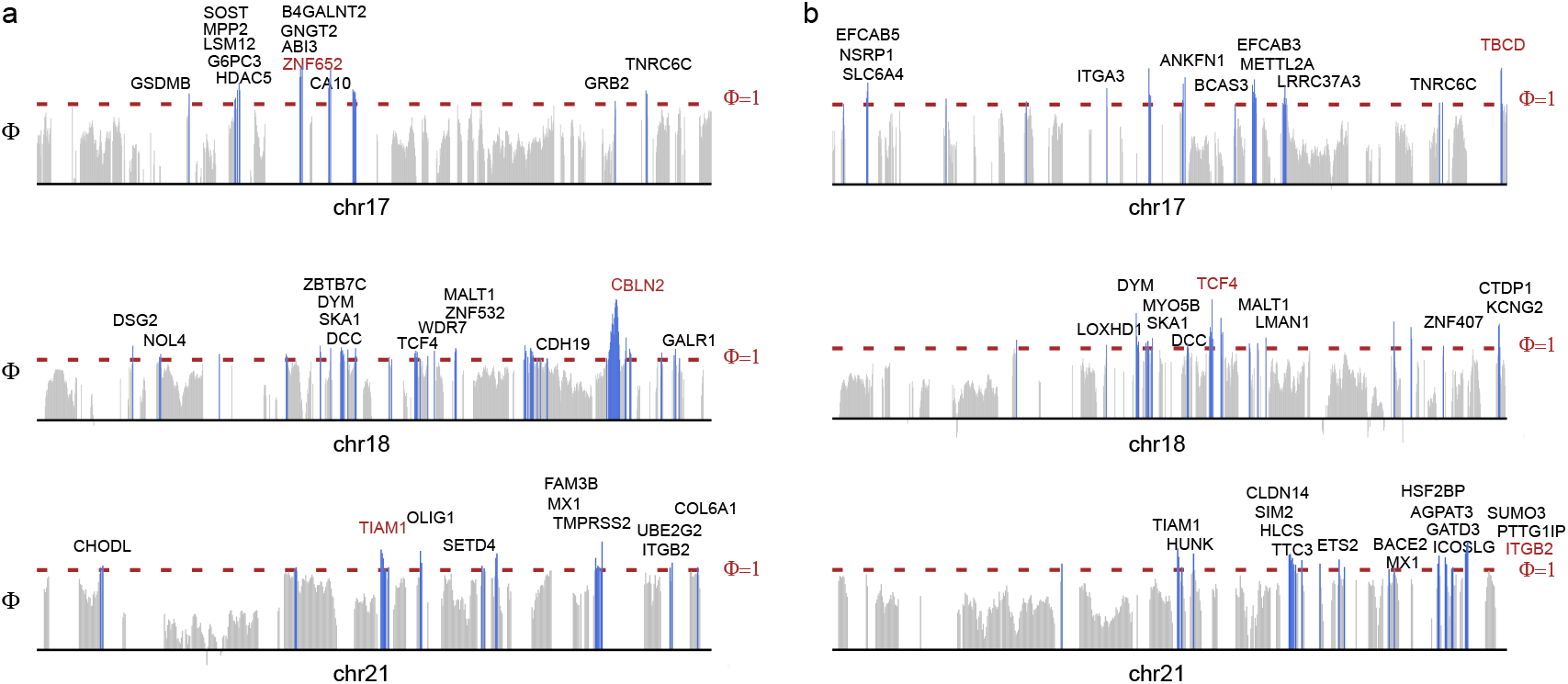
Φ value distribution and the locus–locus network of the lung cancerization (*a*) as well as the reversion process (*b*). The hot spots of lung cancerization are defined as the loci with Φ value higher than 1.0 (colored in blue). The hot spots involved in the protein-coding regions are labeled. The genes with highest Φ value are colored in red.

**Figure 9.**
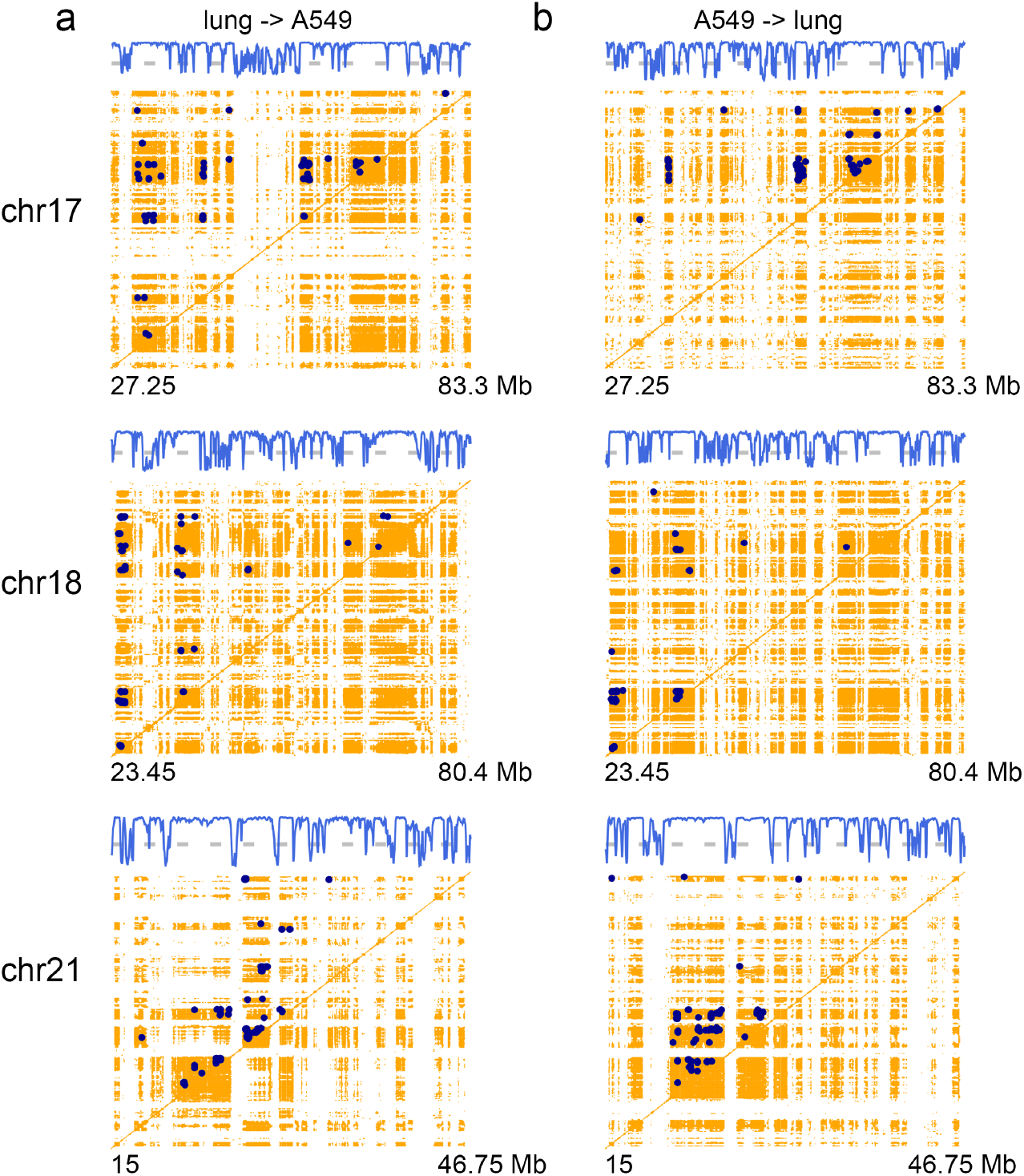
The locus–locus correlation during the lung cancerization (*a*) as well as the reversion process (*b*). The correlation value between locus *i* and locus *j* is quantified through the Pearson correlation coeffcient (*R*^2^) of the local CFI traces of locus *i* and locus *j*. High correlation loci pairs (*R*^2^ *>* 0.7) suggest that locus *i* and locus *j* move synchronously during cancerization or reversion, being illustrated with orange dots. The *“*strong correlation network*” ij* pairs (*R*^2^ *>* 0.95, top 100 *ij* pairs with (*i* − *j*) *>* 10) are labeled with dark-blue points, including coding (genes) and non-coding regions. The locus–locus interaction network can be shown by projecting the cross correlation matrix into one-dimension sequence of the first principal component (pc1, blue lines). The gray dashed lines locate at the zero pc1 value. Here high positive pc1 values correspond to the loci that make up the network.

Of all the genes with high Φ values in this study, we find out that some genes (MPP2, ABI3, GRB2) located in chromosome 17 are related to Src-homology 3 (SH3) domain. The SH3 domain is a famous tumor target that are required for the transmission of proliferative signals initiated by tyrosine kinases ***Smithgall (1995); Gril et al. (2007); Kurochkina and Guha (2013)***. Here these genes have high Φ values in the cancerization pathway, but not in the reversion pathway. Han *et al*. have reported that increased MMP2 expression correlates with malignant biological behavior of lung cancer ***Han et al. (2020)***. The experimental results of Mitra *et al*. indicated that the overexpression of GRB2 can enhance the Epithelial-to-Mesenchymal Transition (EMT) of A549 cells ***Mitra et al. (2018)***. In addition, some genes can influence GTP binding and GTPase activity, including the GNGT2 of chr17, TIAM1 of chr21, MX1 of chr21, SLC6A4 of chr17, MYO5B of chr18. These genes have high Φ values in cancerization or reversion pathway. In general, the aberrant GTP binding regulation or GTPase activity have been observed in many human diseases including many different types of cancer ***Samatar and Poulikakos (2014); Haga and Ridley (2016)***. Previous reports have shown that the TIAM1 is a gene product upregulated in neuroendocrine small-cell lung cancer (SCLC) cells ***Payapilly et al. (2021)***. Some Zinc Finger Protein-coding genes are involved in the lung cancer, such as ZNF652 (chr17), ZNF532 (chr18), and ZNF407 (chr18). The zinc finger containing proteins (ZNFs) that can act as the transcription factors play an important role in many biological processes, including development, differentiation, metabolism and autophagy ***Jen and Wang (2016)***. Besides, we notice that some genes with high Φ values are related to the calcium ion binding or calcium channel activity, including DSG2 (chr18), CDH19 (chr18), EFCAB5 (chr17), EFCAB3 (chr17), and LOXHD1 (chr18). It has been reported that altered expression of specific Ca^2+^ channels and Ca^2+^-binding proteins are characterizing features of lung cancer, which regulate cell signaling pathway leading to cell proliferation or apoptosis ***Yang et al. (2010); Tran (2021)***. The biological analysis has shown that the expression of DSG2 is associated with non-small cell lung cancer (NSCLC) ***Cai et al. (2017)***. Moreover, HDAC5 has a function of histone modification, which displays a significant upregulation in lung cancer ***Zhong et al. (2018)***. The NOL4, which enables RNA binding activity, has been found to be a novel nuclear marker of SCLC cells ***Lee et al. (2022)***. And OLIG1 controls the activity of protein dimerization. The expression of OLIG1 significantly correlates with overall survival in NSCLC patients ***Brena et al. (2007)***. These results demonstrate the crucial role of these genes in the lung cancer. Intriguingly, it should be noted that a large part of these high Φ loci as well as the network components are occupied with non-coding regions. Therefore, the non-coding regions also take a significant part during the transition of lung cancerization or reversion.

A large number of genes belong to the correlation network of lung cancerization or reversion. Here we list the genes involved in the interactions with highest correlation values (*“*strong correlation network*”* with *R*^2^ *>* 0.95, top 100 *ij* pairs). In chromosome 17, the strong correlation network includes NF1, RAB11FIP4, LRRC37B, ZNF207, MYO1D, ASIC2, CCL2, LIG3, KRT, GAST, HIP1, COIL, SCPEP1, RNF126P1, AKAP1, MSI2, CUEDC1, VEZF1, DYNLL2, OR4D1, EPX, MKS1, LPO, BCAS3, AXIN2, CEP112, PRKCA, CACNG5, PITPNC1, NOL1, ARSG, PRKAR1A, FAM20A, ABCA, MAP2K6, SLC39A11, RBFOX3, TBCD (cancerization); MYO1D, AATF, ACACA, RNF126P1, MSI2, CUEDC1, SRSF1, OR4D1, HSF5, RGS9, CEP112, APOH, PRKCA, CACNG4, HELZ, PSMD12, PITPNC1, BPTF, AMZ2, ARSG, PRKAR1A, FAM20A, ABCA, MAP2K6, GPRC5C, CD300, C1QTNF1, RBFOX3, ENPP7 (reversion). In chromosome 18, the strong correlation network includes ZNF521, NOL4, DTNA, MAPRE2, FHOD3, TPGS2, KIAA1328, SLC14A2, CDH20, RNF152, ZCCHC2, CDH7, CDH19, DSEL, DOK6, NETO1 (cancerization); ZNF521, DSC3, DTNA, MAPRE2, ZNF397, FHOD3, TPGS2, KIAA1328, SLC14A2, CDH19, DOK6 (reversion). In chromosome 21, the strong correlation network includes NCAM2, ADAMTS, N6AMT1, GRIK1, KRTAP, MIS18A, DSCAM, PCBP3, COL6A1 (cancerization); NCAM2, ADAMTS, N6AMT1, KRTAP, COL6A1, LSS (reversion). These genes move synchronously during cancerization or reversion process. If one gene was changed, other loci of the network may be influenced. Both the PRKCA and the PRKAR1A genes have been reported to be affected in the melanocytic neoplasm patient samples ***Bahrami et al. (2016)***. RNA-binding proteins, coding by MSI2 (Musashi RNA-binding protein 2) and RBFOX3, have been shown to regulate miRNA biogenesis at the post-transcriptional level ***Michlewski and Cáceres (2019)***. In addition, the works of Nordquist *et al*. have identified that both NCAM2 and N6AMT1 act together for normal pharyngeal pumping and neuromuscular behaviors ***Nordquist et al. (2018)***. The non-coding regions also contribute significantly to the correlation network, which suggests that the genes and non-coding regions can act together during the process from normal lung to lung cancer or the reversed process.

## Conclusion

In this study, we investigate the changes of chromosome structures and functions during the lung cancerization process through the chromosome structural ensemble switching model (CSESM) between lung and A549 cell types. We introduce chromosome ensemble fluctuation quantities, local chromosome fluctuation index (local CFI, CFI_*i*_) and microscopic-level of CFI (micro CFI, CFI_*ij*_), and represent the relationship among CFI quantities, contact probability, epigenetic marks (compartment), and gene expressions. The results show the strong correlations among these chromosome characteristics. In addition, the local CFI has an important role in distinguishing the different chromosomes. Normal lung and lung cancer cell have obvious differences on micro CFI distribution that the cancerization (from normal lung to A549) will increase the relative fluctuation and cause the movement of the large chromosome domain. Moreover, the lung cancer chromosome has more higher degree of the phase separation between compartments A and B. The distribution of local CFI and compartment suggests that the significant *“*switching off*”* events (conformational switching) contribute a lot to the lung cancerization. Similar situations have been observed in the RNA-seq data that the significant gene *“*switching off*”* loci is more than the *“*switching on*”* ones.

The normal lung and lung cancer cell have significant differences on the chromosome structure and function. For the kinetic details, the lung cancerization pathway is not the same as the reversion pathway. We show the kinetic hot spots and networks of lung cancerization by calculating the Φ and correlation values. Simular to proteins, the high Φ loci may determine the mechanism of lung cancer. These hot spots will play an important role during the lung cancer transitions. In addition, the correlation network includes the loci move synchronously during cancerization or reversion process. In summary, the results of different chromosomes indicate the similar trendency of chromosome structural characteristics on the cancerization process. Our theoretical studies elucidate the chromosome structure–function mechanism of lung cancerization, which provide valuable insights on the prevention and regulate of the lung cancer.

## Methods

### Chromosome models and simulation settings

We used the models from Nucleome Data Bank (NDB, https://ndb.rice.edu) ***Contessoto et al. (2021)*** to build the initial structures and potential energies of chromosomes 17, 18, and 21, for different types of human cells (normal lung cell (lung) and lung cancer cell (A549)). The NDB enables physics-based chromosome models for molecular dynamic simulations, by combining the MEGABASE and MiChroM computational pipelines ***Di Pierro et al. (2016***, 2017). The MiChroM model is a 3D chromatin polymer chain at 50 kb resolution, including the epigenetic information from the Encode Project database (epigenetic marking patterns of compartments and sub-compartments from ChIP-seq assays). Therefore, it is a type of block-copolymer (heteropolymer) model with given potential form

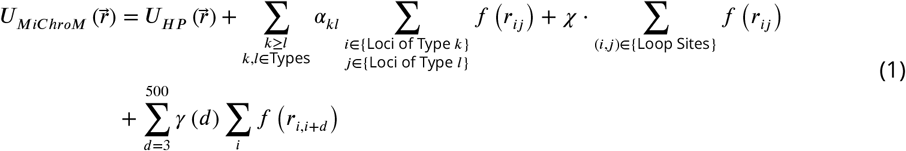

where *U*_*HP*_ is the homo-polymer potential term of the chromosome connectivity, in resolution of 50 Kb of DNA, the second term describes the type–type interactions of the chromosome where the type is the compartment annotation determined by MEGABASE, the third term is the interactions between loop anchors that are related to the protein CTCF, the final term is the one referred to the ideal chromosome model. Further details for this model can be referred to the reference of NDB ***Contessoto et al. (2021)***. This model is transferable for different chromosomes and cells, and has been shown to successfully predict the Hi-C contact maps of multiple human cell lines, and is consistent with the fluorescence in situ hybridization (FISH) experimental results ***Contessoto et al. (2021); Cheng et al. (2020); Junior et al. (2021)***.

Here in this study, the 1666-, 1608-, and 935-bead models (*n* = 1666, *n* = 1608 and *n* = 935), representing the human chromosomes 17, 18, and 21 were prepared for molecular dynamics (MD) simulations, respectively. The simulations were performed by Gromacs 2018 ***Abraham et al. (2015)*** with reduced units. Langevin stochastic dynamics were applied with a friction coefficient of 1.0 *τ* ^−1^, where *τ* is the reduced time unit. In order to obtain the ensemble-average properties of the chromosomes, such as the probability contact maps and the fluctuations, we constructed the ensemble of chromosome after a two-stage simulation process ***Zhang and Wolynes (2016)***: a heating stage (2 × 10^5^ *τ* at temperature 3.0 in reduced time unit) to relax the initial model inside the nucleus wall as well as a replica-exchange stage (1 × 10^5^ *τ* of 28 replicas, all at about 1.0 temperature in reduced time unit) to enhance the sampling of the chromosome structures. Then an ensemble of 2800 chromosome frames were collected from the last 5 × 10^4^ *τ* of the 28 replicas (100 frames each).

We developed the chromosome structural ensemble switching model (CSESM), which is similar as the previous energy landscape-switching model ***Chu and Wang (2020a***,b, 2021), encoding the epigenetic information to perform the cancerization and reversion processes from lung to A549 and from A549 to lung. From a physical perspective, the energy landscapes of two cell types of the cancerization/reversion will differ significantly, form structural to energy. The energy excitation– relaxation landscape-switching model is developed for the non-equilibrium process to connect the two distinct energy landscapes. The non-equilibrium effects in the landscape-switching model are the driving forces for the cancerization/reversion processes, which is often in the form of extended ATP hydrolysis for energy pumping in biology. As the previous simulation procedures ***Chu and Wang (2020a***,b, 2021), here the CSESM simulations were performed under the potential of the beginning state of cancerization/reversion. Then the potential was suddenly switched to that of the end state of cancerization/reversion. Finally, simulations were performed under the potential of the end state of cancerization/reversion. Considering the ensemble of the chromosome, 2800 runs were performed for each cancerization/reversion process. For each process, 2.0 × 10^3^ *τ* is long enough for one type of chromosome ensemble to transform to another. We can obtain the important information on the chromosome ensemble at each time step.

### Statistical analysis

For each chromosome ensemble, the probability contact map *P* can be calculated as 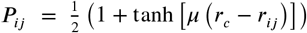, where *μ* = 3.22 and *r*_*c*_ = 1.78 were set according to the previous studies ***Zhang and Wolynes (2016); Di Pierro et al. (2016***, 2017); ***Contessoto et al. (2021)***. According to the probability contact map *P*, the A and B compartments were identified via the same method used by Lieberman-Aiden *et al*. ***Lieberman-Aiden et al. (2009); Nagano et al. (2013); Imakaev et al. (2012)***. That is, the enhanced probability contact map, calculated by the observed/expected matrix, was normalized by ICE method and then converted into a Pearson correlation matrix. The compartment profiles were determined by the first principal component (pc1) of the principal component analysis (PCA) of the matrix ***Abdi and Williams (2010)***. Within each ensemble, the relative flexibility of a contact pair between *i* and *j* was measured via a dimensionless quantity 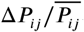, where the Δ *P*_*ij*_ and 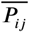 are the standard deviation and the mean of all the *P*_*ij*_ among the chromosome ensemble. Then we use the local chromosome flexibility index (local CFI) to quantify the degree of the fluctuations of the chromosome at position *i* in the ensemble: 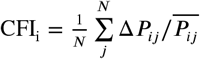. The CFI spectrum (CFI_i_ values of the chromosome) quantifies the fluctuation distribution of the chromosome. Micro CFI 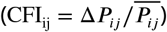 is defined to quantify the degree of the fluctuations of a pair of loci *i* and *j* in the ensemble.

In this study, we determine a pair of reaction coordinates of compartment: *κ*_*A*_ (*α* ln(0.05 − pc1_*i*_)(0.05 − pc1_*j*_)) and *κ*_*B*_ (*α* ln(0.05 + pc1_*i*_)(0.05 + pc1_*j*_)). Here the pc1_*i*_ and pc1_*j*_ are the first principal component (pc1) values of loci *i* and *j*. When *i* and *j* belong to the same compartment, (A–A or B–B, both pc1_*i*_ and pc1_*j*_ larger than 0 or lower than 0), *α* = 1. When *i* and *j* belong to different compartments, (A–B, both pc1_*i*_ × pc1_*j*_ lower than 0), *α* = −1. As a result, the interactions within compartment (A–A and B–B) and between different compartments (A–B) have different data ranges. In addition, for better comparison of different compartments, both *κ*_*A*_ and *κ*_*B*_ were calculated for analysis.

### RNA-seq and DNA methylation data

All the RNA-seq (total RNA-seq) data were downloaded from ENCODE ***Consortium et al. (2012)***, including the lung (ENCSR917YHC) and A549 (ENCSR000CTM). All the DNA methylation (WGBS) data were obtained from ENCODE ***Consortium et al. (2012)***, including the lung (ENCSR556KEJ) and A549 (ENCSR481JIW). Both plus and minus strands signal of all reads were used for comparison.

## Acknowledgments

W.-T.C. thanks Network and Computing Center, Changchun Institute of Applied Chemistry, Chinese Academy of Science and Computing Center of Jilin Province for computational support. W.-T.C. thanks the support from National Natural Science Foundation of China Grants (32171244, 12234019, 21721003), Youth Innovation Promotion Association CAS Grant (2020231), and Jilin Province Science and Technology Development Plan Grant (20200301009RQ).

