## Supplementary material for "Uncovering the lung cancer mechanisms through the chromosome structural ensemble characteristics": Figure S1

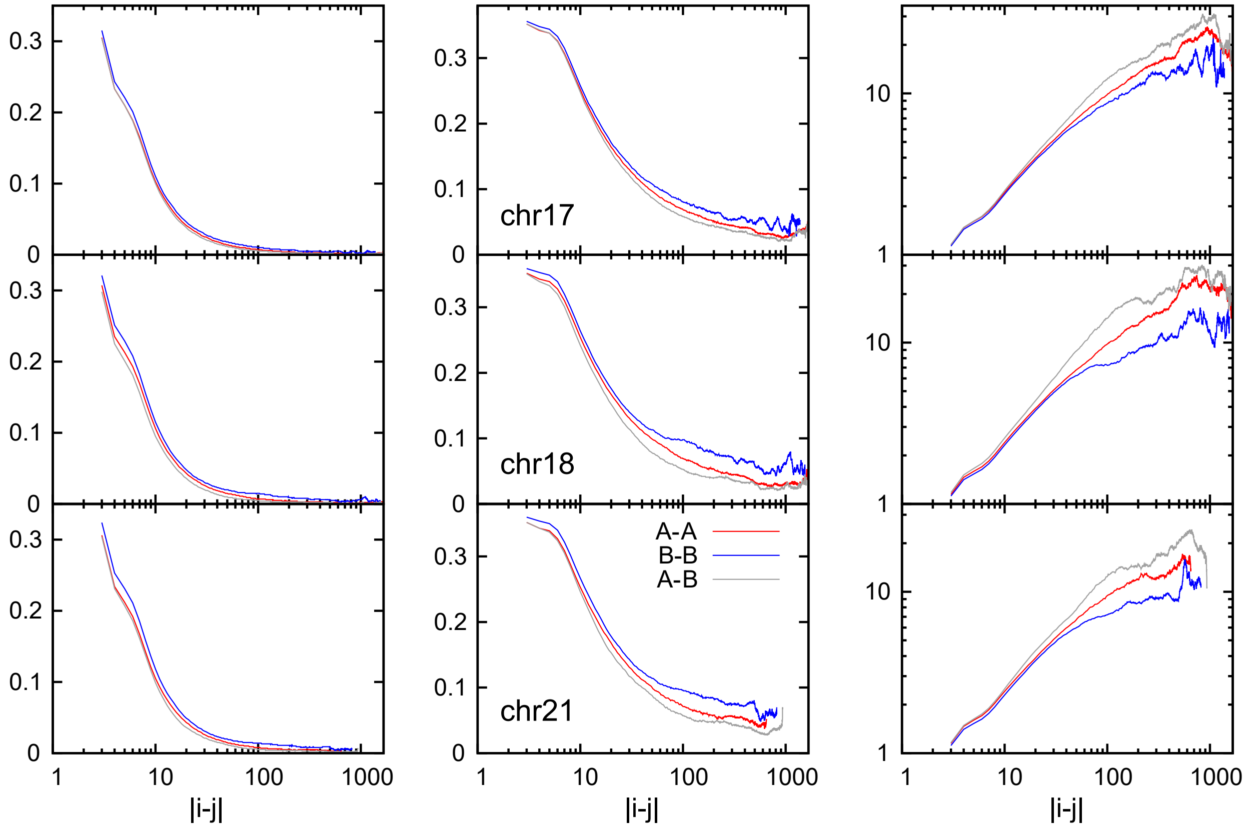


Figure S1: The mean contact probability $\overline{P_{ij}}$ (left), the standard deviation $\Delta P_{ij}$ (middle), and the ratio of them $\Delta P_{ij}/\overline{P_{ij}}$ (micro CFI, right) within the compartment A (A–A), within the compartment B (B–B), and between compartment A and B (A–B) as a function of the chain distance $|i-j|$ in the normal lung cell. Curves of chromosome 17, 18, and 21 are labeled in this figure.

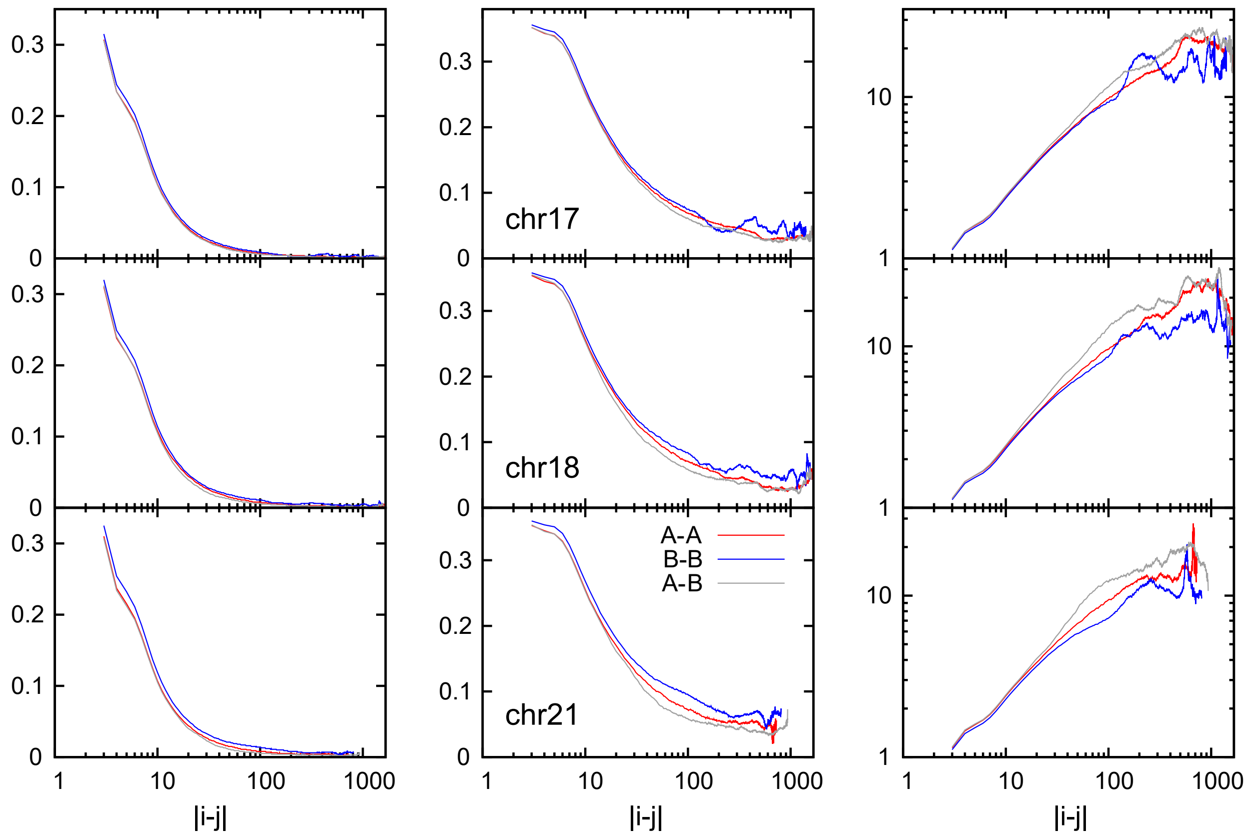


Figure S2: The mean contact probability $\overline{P_{ij}}$ (left), the standard deviation $\Delta P_{ij}$ (middle), and the ratio of them $\Delta P_{ij}/\overline{P_{ij}}$ (micro CFI, right) within the compartment A (A–A), within the compartment B (B–B), and between compartment A and B (A–B) as a function of the chain distance $|i-j|$ in the lung cancer A549 cell. Curves of chromosome 17, 18, and 21 are labeled in this figure.

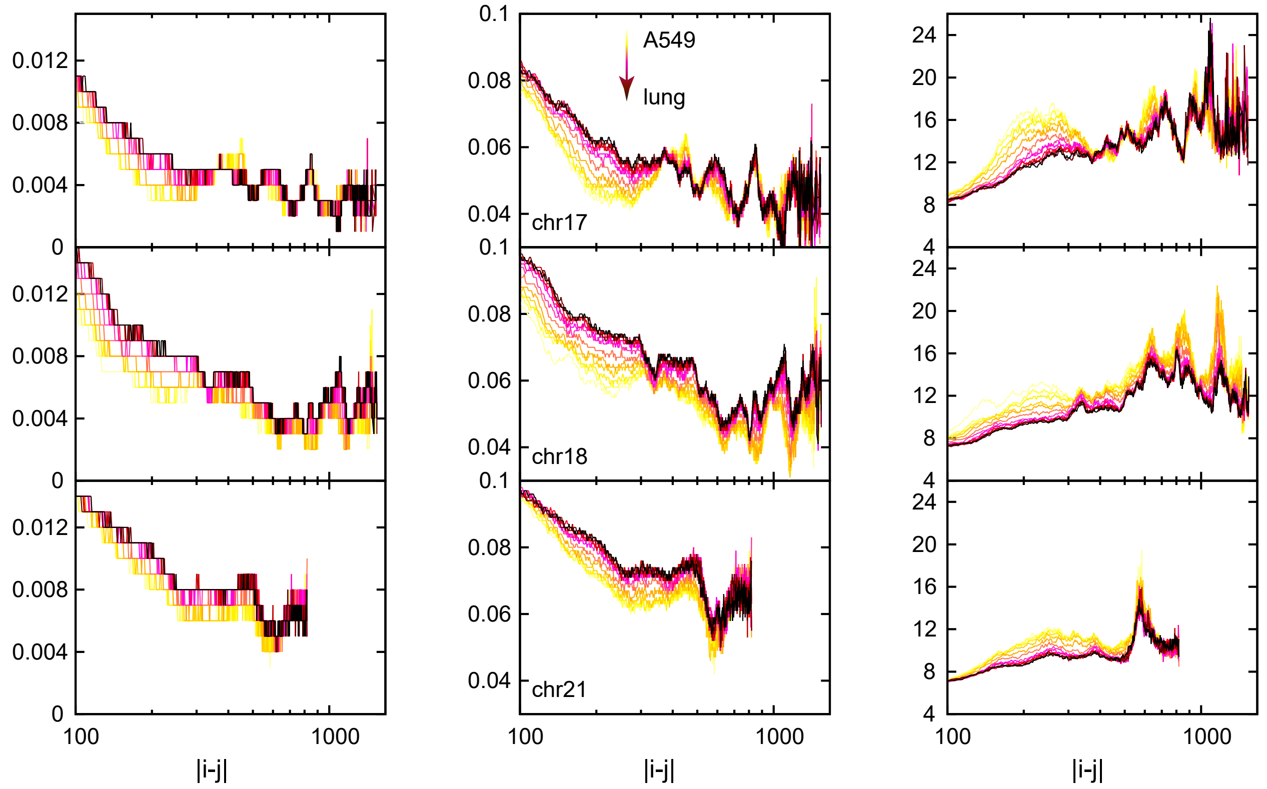


Figure S3: The mean contact probability $\overline{P_{ij}}$ (left), the standard deviation $\Delta P_{ij}$ (middle), and the micro CFI (right) within the compartment B (B–B) as a function of the chain distance $|i-j|$ during the pathway from the lung cancer A549 cell to the normal lung cell. Curves of chromosome 17, 18, and 21 are labeled in this figure.

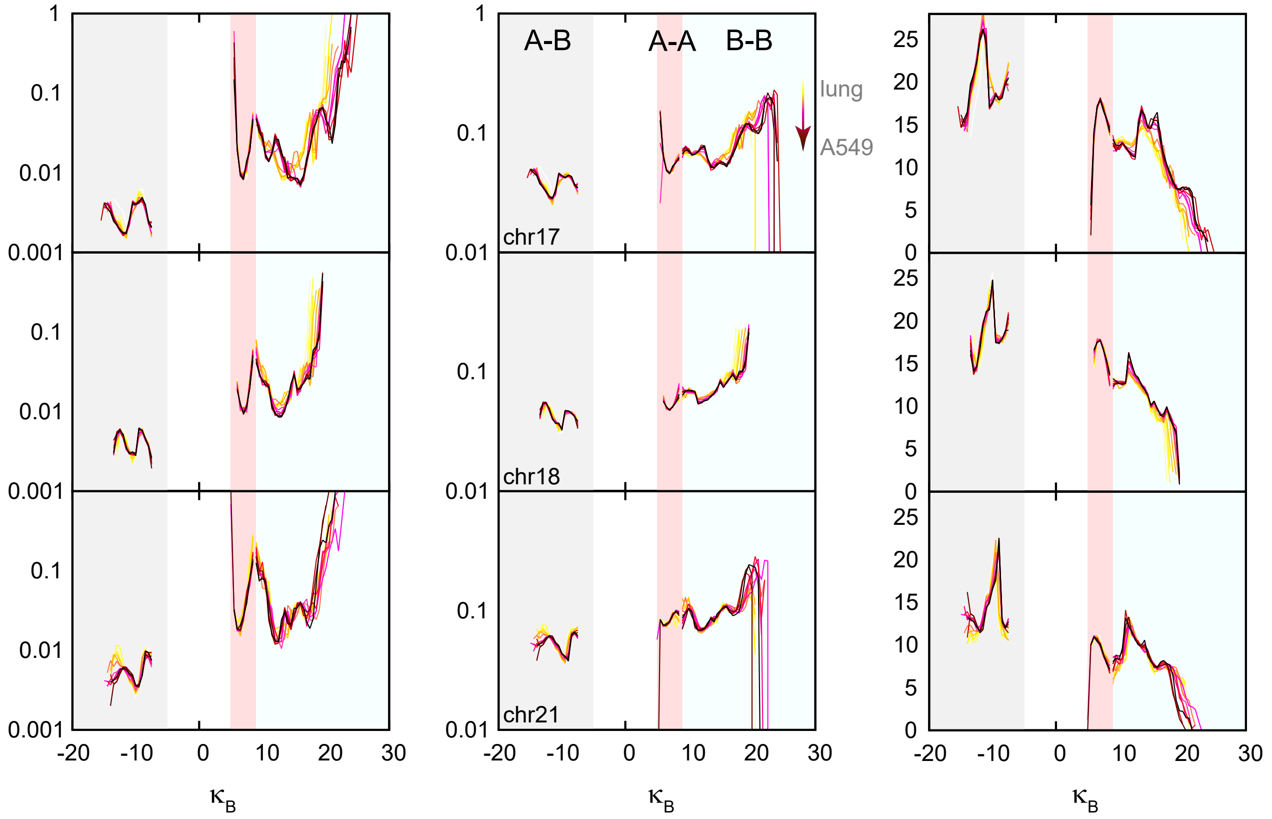


Figure S4: The distribution between $\kappa_B$ ($\alpha \ln(0.05 + \mathrm{pc1}_i)(0.05 + \mathrm{pc1}_j)$) and $\overline{P_{ij}}$ (left), between $\kappa_B$ and $\Delta P_{ij}$ (middle), between $\kappa_B$ and micro CFI during the pathway from the normal lung cell to the lung cancer A549 cell. The regions of different compartments, the curves of chromosome 17, 18, and 21 are labeled in this figure.

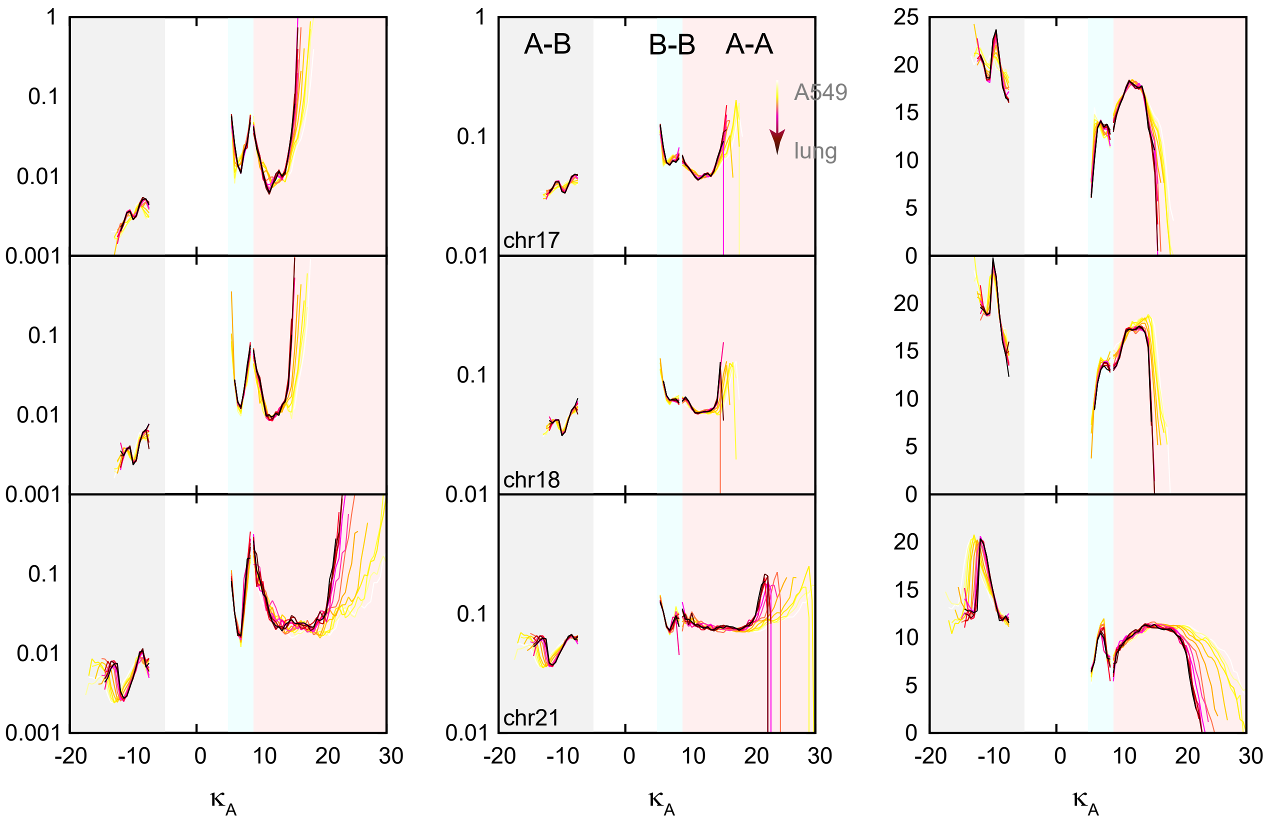


Figure S5: The distribution between $\kappa_A$ ($\alpha \ln(0.05 - \text{pc1}_i)(0.05 - \text{pc1}_j)$) and $\overline{P_{ij}}$ (left), between $\kappa_A$ and $\Delta P_{ij}$ (middle), between $\kappa_A$ and micro CFI during the pathway from the lung cancer A549 cell to the normal lung cell. The regions of different compartments, the curves of chromosome 17, 18, and 21 are labeled in this figure.

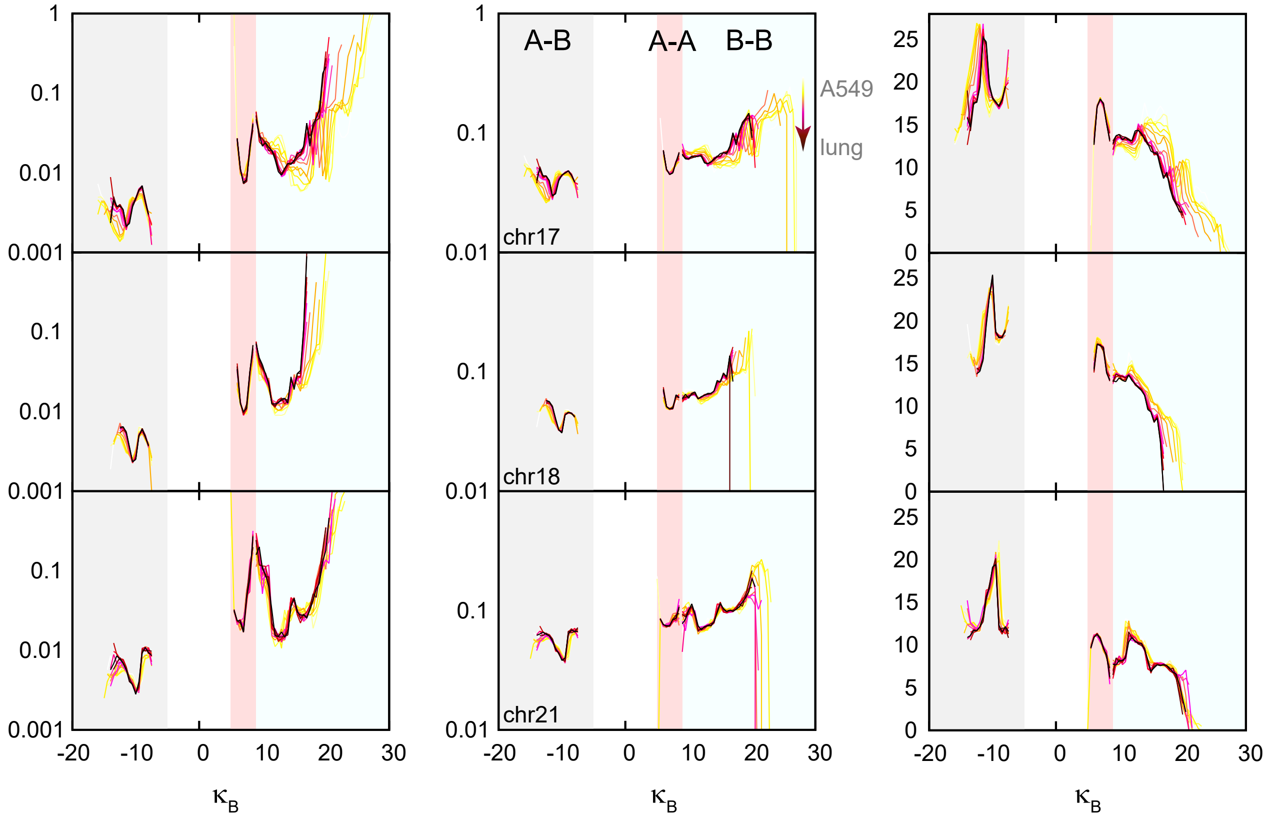


Figure S6: The distribution between $\kappa_B$ ($\alpha \ln(0.05+\text{pc1}_i)(0.05+\text{pc1}_j)$) and $\overline{P_{ij}}$ (left), between $\kappa_B$ and $\Delta P_{ij}$ (middle), between $\kappa_B$ and micro CFI during the pathway from the lung cancer A549 cell to the normal lung cell. The regions of different compartments, the curves of chromosome 17, 18, and 21 are labeled in this figure.

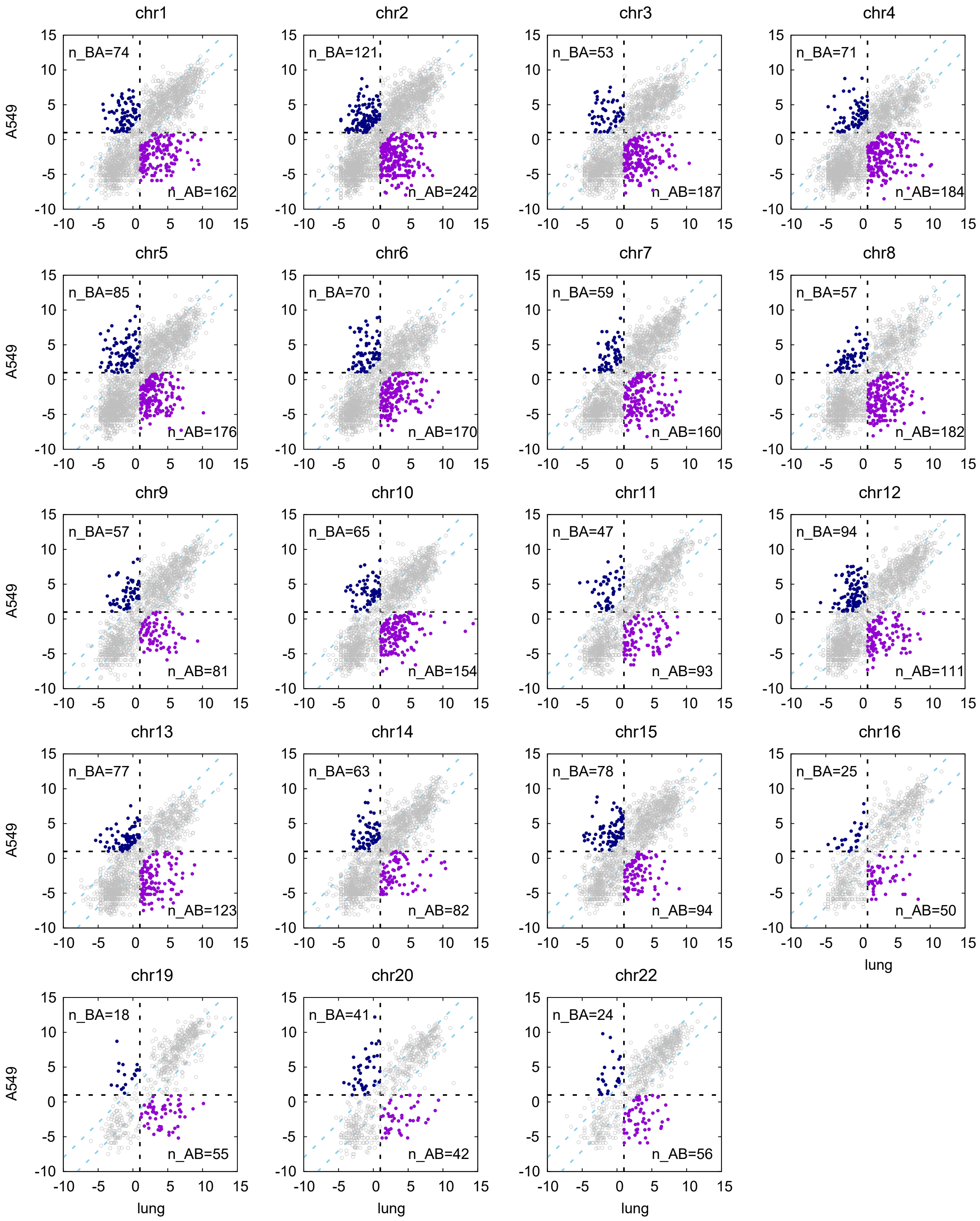


Figure S7: The different gene expression level (quantified with ln(RNA-seq signal)) of each locus (in 50 kb resolution, *q* arm) in the normal lung and the lung cancer (A549) cells. Here the data of other chromosomes except for the 17, 18, and 21 are listed in this figure. The significant gene "switching off" (A–B, purple dots) or "switching on" (B–A, blue dots) loci during the cancerization process are defined the same as those in Figure 7. The number of gene "switching off" (n_AB) and "switching on" (n_BA) loci is labeled in this figure.

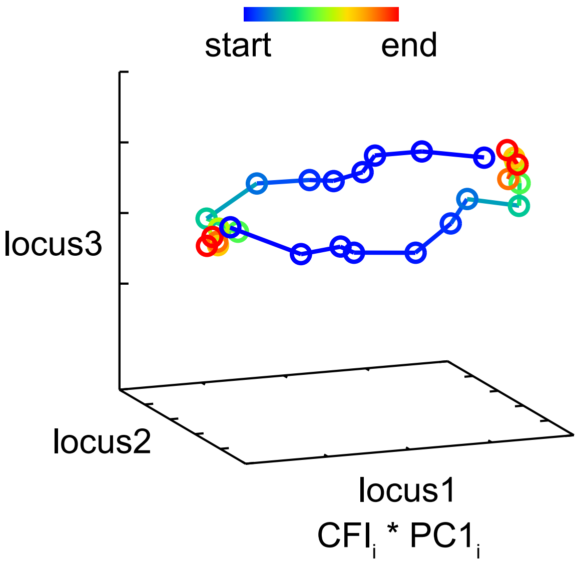


Figure S8: The the forward trace and the reverse trace projecting on three loci. The reaction coordinate is defined as the $\mathrm{CFI}_i \times \mathrm{pc1}_i$ of locus $i$. The trace curve is colored with the kinetic time.
